# A Proof-of-Principle of Nanoscale Optogenetics

**DOI:** 10.1101/395475

**Authors:** Markus A. Stahlberg, Charu Ramakrishnan, Katrin I. Willig, Edward S. Boyden, Karl Deisseroth, Camin Dean

## Abstract

Optogenetics has revolutionized the study of circuit function in the brain, by allowing activation of specific ensembles of neurons by light. However, this technique has not yet been exploited extensively at the subcellular level. Here we propose a novel focal stimulation approach using STED/RESOLFT-like illumination, whereby switchable light-gated channels are focally activated by a laser beam of one wavelength and deactivated by an overlapping donut-shaped beam of a different wavelength, confining activation to a center focal region. We demonstrate the utility of current optogenetic tools to achieve highly focal depolarization using this method and further examine a proof-of-principle of nanoscale optogenetic activation using an initial macroscale approach. When employed at the nanoscale, this approach will allow unprecedented optogenetic control of nanodomains within cells.

## Introduction

Approaches to examine the biological mechanisms underlying synaptic plasticity rely on the artificial stimulation of neurons. Since its discovery, channelrhodopsin 2 (ChR2) - a channel that depolarizes neurons when activated by blue light - and its derivatives, have become the dominant technique used to activate or inactivate neurons (Deisseroth, 2011). So far, the stimulation of neurons using channelrhodopsins has mainly focused on targeting specific cell populations or areas in the brain of rodents (Gradinaru et al., 2010; Gunaydin et al., 2010). But the ability to stimulate single synapses - ranging from 0.04 *μ*m^2^(Schikorski and Stevens, 1997) to approximately 0.1 *μ*m^3^ for a dendritic spine - or even sub-synaptic nanodomains - using channelrhodopsins, would be particularly advantageous to study aspects of synapse-specificity and plasticity, for example.

Several attempts have been made to decrease the area of activation of channelrhodopsins. Spatially controlled stimulation of postsynaptic sites has been achieved by optogenetic stimulation of the presynaptic neuron (Zhang and Oertner, 2007), for example. An activated area less than 30 μm was achieved by focused laser beams with reduced laser power (Schoenenberger et al., 2008), and two-photon stimulation with 920 nm light has been used to evoke action potentials in individual cells (Mohanty et al., 2008; Packer et al., 2012; Papagiakoumou et al., 2010). However, so far no technique exists that confines stimulation of light-gated channels to microscale - or even nanoscale - subcellular regions, to stimulate individual synapses or sub-synaptic sites. We developed such an approach by making use of mutations in light-gated channels that slow the photocycle, making it easier to control their switching between active and inactive states in response to two different wavelengths of light.

The photocycle of ChR2 has been intensively studied (Bamann et al., 2008, 2010; Nikolic et al., 2009; Stehfest et al., 2010). When ChR2 is illuminated with blue light, it undergoes a conformational change from its dark-adapted closed state (D470), through two non-conducting intermediate states (P500 and P390) and enters the open channel state (P520). Subsequently, it passes through several intermediate states, until it finally reaches its dark-adapted closed state again and the cycle can be repeated. Interestingly, the P390 intermediate state, and the P520 open state, are sensitive to UV light (∼390 nm) and green light (∼520 nm), respectively, which mediate a direct transition to the closed channel-state (D470) (Stehfest et al., 2010). ChR2 channels that possess a mutation at residue C128 - a conserved amino acid that interacts with the all-trans retinal Schiff base chromophore - have delayed inactivation and a prolonged open state following blue light activation, which can be converted directly to the closed state by green light, creating a so-called “step function” opsin (Yizhar et al., 2011). The ChR2 C128S, C128T and C128A step-function opsin mutants, possess long open-channel states. These channels can be efficiently closed by exposure to a second closing wavelength (and opened again by an activating wavelength). However, the mutations that cause channels to close in response to a second wavelength of light, often result in a dramatic reduction in photocurrent, at least for existing channels with these switchable mutations (Yizhar et al., 2011).

Here we tested a focal activation approach using step function opsin variants with larger photocurrents, and an illumination technique inspired by STED/RESOLFT microscopy: A center focused light beam is used to activate channels, while a surrounding overlapping donut illumination of the respective closing wavelength of channels eliminates the photocurrent in this area, resulting in a highly focal site of activation. This strategy, if employed in sub-diffraction limited regions, would allow unprecedented nanoscale focal activation. We demonstrate the utility of current optogenetic tools to achieve highly focal depolarization using this method, and further examine a proof-of-principle of nanoscale optogenetic activation, using an initial macroscale approach.

## Results

Focal stimulation of channelrhodopsins requires that activated channels remain relatively immobile in cell membranes. We therefore first tested the diffusion of channelrhodopsins expressed in dissociated hippocampal neurons. STED microscopy images of transfected dissociated hippocampal neurons revealed that ChR2-EYFP was mainly targeted to the plasma membrane of neurons (**Fig. 1A**). To examine channel diffusion, we performed fluorescence recovery after photobleaching (FRAP) experiments. A 2.5 *μ*m region was illuminated with 488 nm light at 100% laser power for one second to bleach ChR2-EYFP in the illuminated region in neuronal processes or cell bodies and imaged for 200 s post-bleaching (**Fig. 1B**). ChR2-EYFP fluorescence in the bleached region recovered partially, with a diffusion coefficient of 3.3 ± 0.2 x 10^−2^ *μ*m^2^/s in processes and 4.7 ± 0.02 x 10^−2^ *μ*m^2^/s in cell bodies, indicating low mobility, such that centrally activated channels would be inactivated by overlapping donut co-illumination 500 nm in width or greater, assuming complete inactivation of step function opsin channels within 1s (Yizhar et al., 2011).

**Figure 1:**
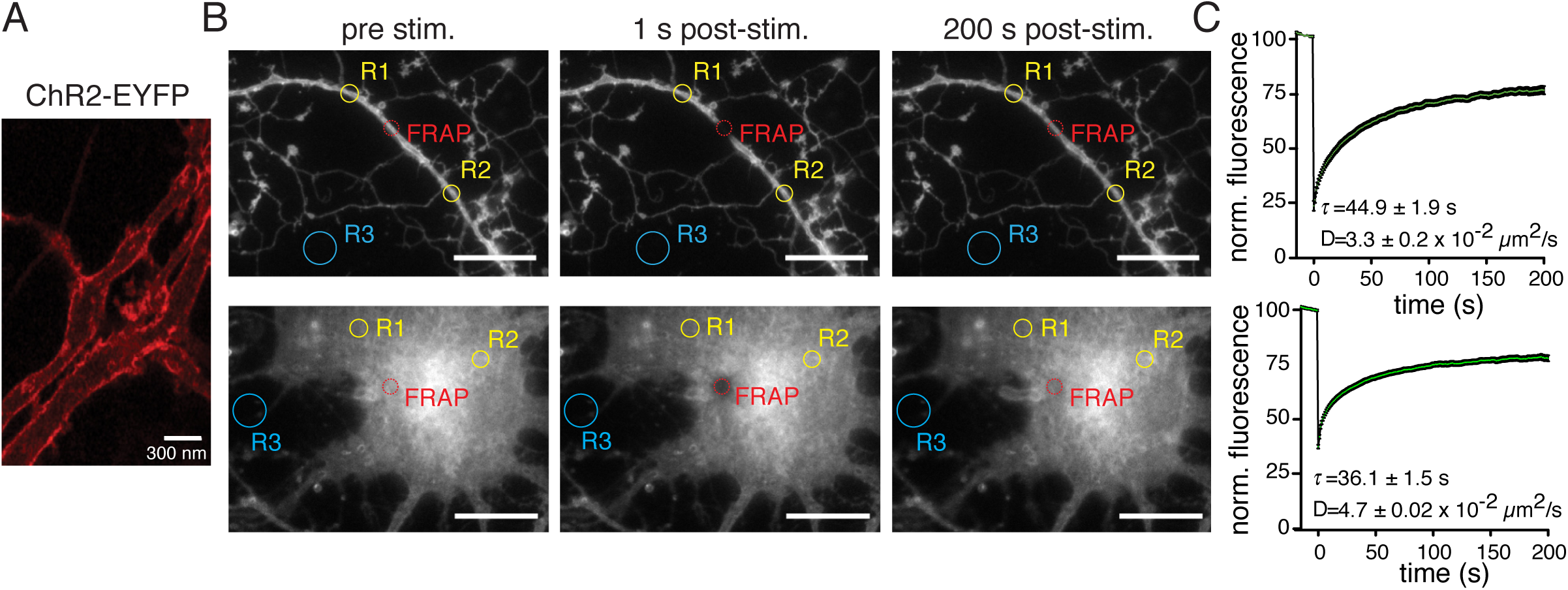
Channelrhodopsin localization and diffusion in the membrane. **A)** STED microscopy image of ChR2-EYFP in the dendrites of a transfected hippocampal neuron in culture. Scale bar = 300 nm. **B)** FRAP experiments of ChR2-EYFP in transfected neuronal processes (top) and cell bodies (bottom) before (left), immediately after (middle) and 200 s after (right) photobleaching of the indicated area. Scale bars = 20 *μ*m. R1 and R2 indicate non-bleached control regions, and R3 indicates background fluorescence black level, used for normalization of fluorescence recovery following photobleaching **(C)**.

A prerequisite for a potential micro/nanoscale focal illumination approach is that channels 1) have a relatively high photocurrent, to allow detection of focal activation, and 2) can be activated by one wavelength of light, and inactivated by simultaneous overlapping illumination with another wavelength. Because high photocurrents, and inactivation of channelrhodopsins with a second overlapping wavelength of light is difficult to predict based on structure-function analysis, we tested a broad range of channelrhodopsin variants for these properties. Dissociated hippocampal cultures were transfected with expression constructs of EYFP or mCherry tagged light-gated ion-channels using calcium phosphate. Channel-specific light evoked photocurrents were then tested to determine channel activation spectra, conductance, gating kinetics and the ability to close channels with a second wavelength, using patch-clamp recordings of cells identified by their respective fluorescence. Spontaneous activity was suppressed by application of TTX, APV, CNQX and gabazine during recordings. For each channel, we measured photocurrents elicited by 405, 488, 561, 594 and 639 nm single wavelength laser illumination. Subsequently, channel inactivation was tested by coillumination with all available laser combinations. Only the wavelength combinations for which effects were observed, are shown in subsequent figures.

The first group of tested channels included the ChR2 variants with comparatively fast-photocycles: H134R, T159C, E123T/T159C and L132C. Fast photocycle channels do not possess a stable open channel state and will open, close and re-open in relatively fast succession. Upon initial exposure to their excitation wavelength all channels open at once and as they proceed through their photocycles, will finally reach an equilibrium (the stationary current phase) of constantly opening and closing channels, in which approximately 60% of all illuminated channels are in the open-channel state (Feldbauer et al., 2009), inferred by the peak to stationary current ratios. The stationary current is very robust; the individual contribution of a single channel is small and of short persistence, because each individual channel closes quickly and re-opens again. Therefore, illumination with a closing wavelength, exciting the conducting P520 or respective state, accelerating channel closure, is thought to have only a minor influence on the total photocurrent of fast photocycle channels. As expected, photocurrents of fast photocycle variants were generally strong and disappeared quickly when illumination ceased (**Fig. 2**).

**Figure 2:**
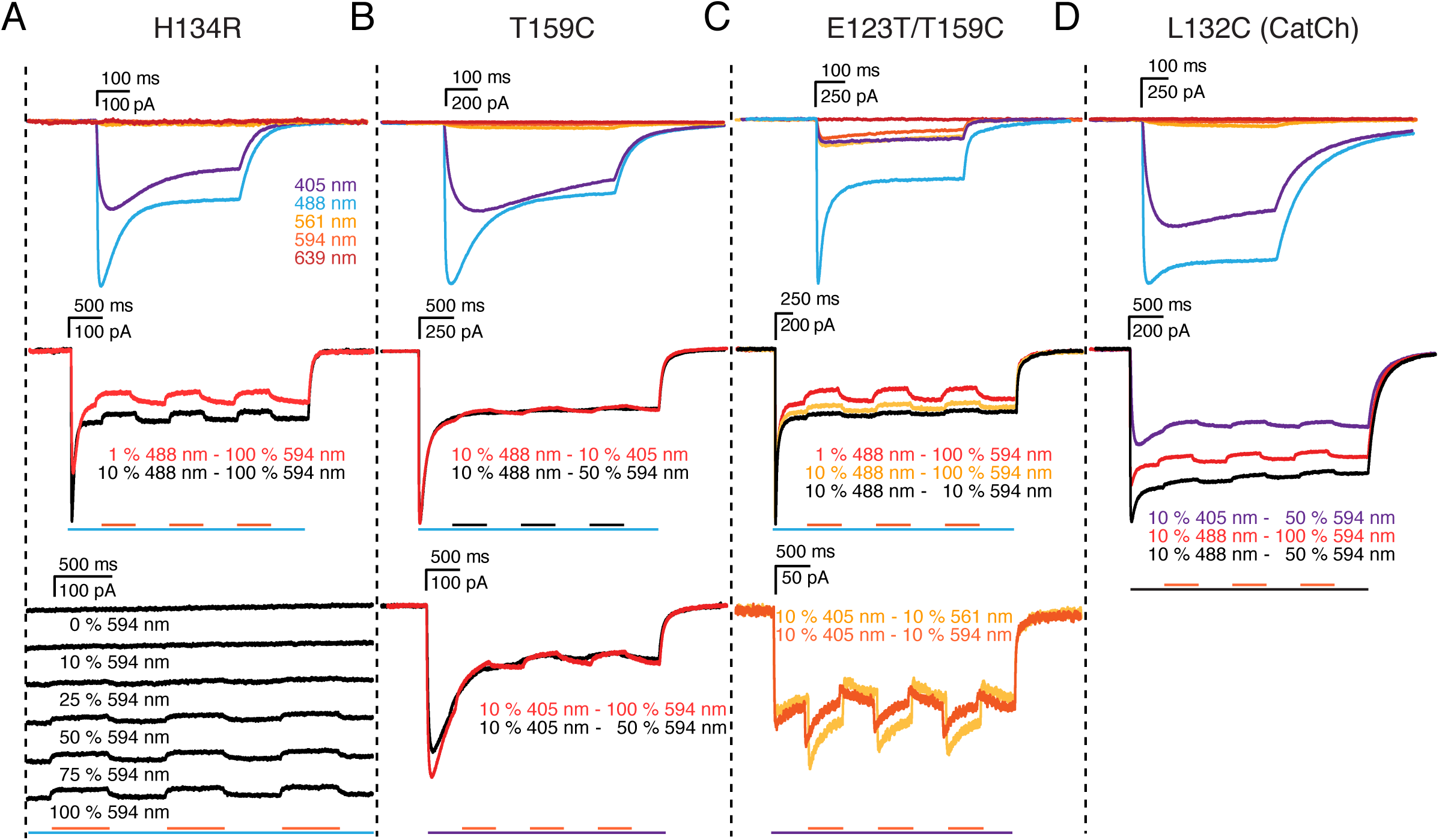
Illumination evoked photocurrents of light-gated channels with fast photocycles. **A**) Photocurrents from recorded hippocampal neurons in culture expressing H134R, T159C (**B**), E123T/T159C (**C**) and L132C (**D**) channels, elicited by 405, 488, 561, 594 and 639 nm single wavelength laser illumination (top panels) and by co-illumination with different wavelengths (lower panels). Photocurrent recordings at specific illumination wavelengths are indicated by the respective color of the recording. Scale bars above current traces indicate recorded current (vertical) and time (horizontal). For coillumination experiments, bars below recordings indicate time of illumination for the indicated color-coded wavelength. Black bars refer to coillumination with the color of the depicted traces in cases where more than one coillumination wavelength was tested. Activating laser intensities were 405 nm 10% (86.83 mW/cm^2^); 488 nm 1% (79.66 mW/cm^2^) and 10% (295.6 mW/cm^2^); 561 nm 10% (146.3 mW/cm^2^) and 639 nm 10% (22.35 mW/cm^2^), and inactivating laser intensities were 594 nm 10% (80.1 mW/cm^2^), 50% (40.06 mW/cm^2^) and 100% (801.48 mW/cm^2^).

The most commonly used fast-photocyle channelrhodopsin variant contains the H134R mutation, which promotes enhanced stationary photocurrents (Nagel et al., 2005). This channel was activated by 405 and 488 nm light, yielding photocurrents of several hundred pA, and was not activated by wavelengths greater than 561 nm (**Fig. 2A, top**). 594 nm light was previously reported to be the most efficient wavelength to close step function ChR2 channels (Yizhar et al., 2011). After opening H134R channels with 488 nm light, overlapping illumination with 594 nm light was the only wavelength that reduced the photocurrent, but only slightly when 10% activating laser power was used, and slightly more when only 1% activating laser power was used (although this reduction in activating laser power also reduced the initial photocurrent) (**Fig. 2A, middle**). Increasing 594 nm laser power increased inactivation accordingly (**Fig. 2A, bottom**).

T159C mutants are reported to possess increased photocurrents (Berndt et al., 2011). Photocurrents were induced by 405 nm and 488 nm light (and not further red-shifted wavelengths) and were indeed approximately twice as large as those of H134R channels (**Fig. 2B, top**). However, these channels showed barely perceptible inactivation with overlapping 405 nm or 594 nm illumination, when activated by either 488 nm light (**Fig. 2B, middle**), or 405 nm light (**Fig. 2B, bottom**). Interestingly, addition of the E123T mutation - which speeds up the photocycle and channel kinetics for temporal precision (Berndt et al., 2011) - further increased peak and stationary currents to more than 1.5 nA and 700 pA, respectively, when channels were activated with 488 nm light, and resulted in weak photocurrents of 250 pA when activated with 405, 561 and 594 nm light. This channel was inactivated slightly by 594 nm light following activation with 488 nm light (**Fig. 2C, middle**). Due to its increased activation in the red spectrum, photocurrents of E123T/T159C elicited by 405 nm light increased even further with simultaneous illumination with 561 nm or 594 nm light (**Fig. 2C, bottom**).

The L132C ChR2 mutant also has high photocurrents and was reported to make channels calcium permeable (Kleinlogel et al., 2011), which could be advantageous to assay focal activation by calcium imaging. This channel displayed strong photocurrents of 750 pA in response to 405 nm activation, and 1.3 nA for 488 nm activation, with a reduced peak-stationary current ratio, compared to other channels. A pronounced slope in decay time indicates slow channel kinetics (**Fig. 2D, top**). However, this channel also exhibited minimal inactivation with 594 nm light following activation with 405 or 488 nm light (**Fig. 2D, bottom**).

The C1V1 and ReaChR red-shifted channels and the Chronos and CoChR channels also have a fast photocycle, but it has not been as extensively studied. The C1V1 channel - comprised of Chlamydomonas and Volvox ChR1 domains, and containing the E122T/E162T mutations that speed up the photocycle (Yizhar et al., 2011) - and ReaChR (Lin et al., 2013) possess an extended activation spectrum towards the red-shifted wavelengths, but responded almost equally well to 405 and 488 nm light and had maximal photocurrents of ∼600 pA (**Fig. 3A, B**). These channels showed little inactivation under coillumination. Photocurrents of C1V1 E122T/E162T activated with 405 nm light were actually further increased by coillumination with 488 nm or 561 nm light (**Fig. 3A**). CoChR and Chronos channels (Klapoetke et al., 2014) could be efficiently opened by illumination at 405, 488 and 561 nm light, where 488 nm evoked the strongest photocurrents (**Fig. 3C, D**); CoChR in particular, featured remarkably strong photocurrents of 3 - 4 nA. CoChR photocurrents could be reduced slightly when 405 nm activation was combined with 561 nm coillumination, and when 488 or 561 nm activation was combined with 405 nm coillumination. This was particularly surprising in the case of 561 nm evoked photocurrents, because in single wavelength tests, 405 nm evoked currents were stronger than those evoked by 561 nm light (**Fig. 3C**). 488 nm coillumination increased photocurrents of the 561 nm activated state of CoChR. Chronos only showed a weak photocurrent reduction for simultaneous 405 nm illumination, when activated with 488 nm light (**Fig. 3D**). On the contrary (and as expected from single wavelength results), 405, 488 or 561 nm coillumination of Chronos transfected neurons increased photocurrents of the 405 or 561 nm activated states.

**Figure 3:**
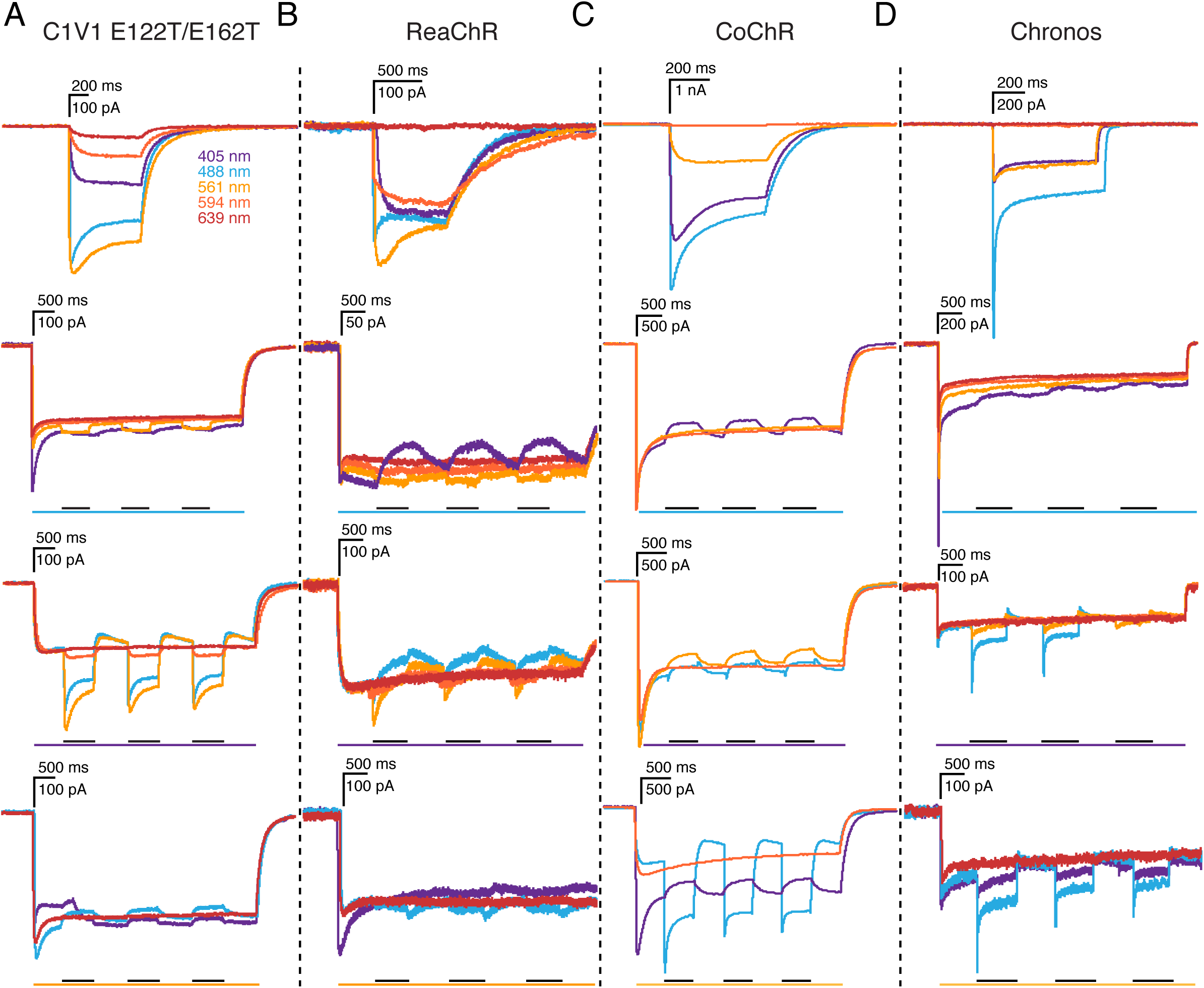
Illumination evoked photocurrents of red-shifted light-gated channels. A-D) Photocurrents of the indicated channels elicited by 405, 488, 561, 594 and 639 nm single wavelength laser illumination (top panels) and by co-illumination with different wavelengths (lower panels). Photocurrent recordings at specific illumination wavelengths are indicated by the respective color of the recording. Scale bars above individual experiments indicate recorded current (vertical) and time (horizontal). For coillumination experiments, bars below recordings indicate time of illumination for the indicated color-coded wavelength. Black bars refer to coillumination with the color of the depicted traces in cases where more than one coillumination wavelength was tested. Activating laser intensities were 405 nm 10% (86.83 mW/cm^2^); 488 nm 10% (295.6 mW/cm^2^); 51 nm 10% (146.3 mW/cm^2^) and 639 nm 10% (22.35 mW/cm^2^) and inactivating laser intensities were 405 nm 100% (662.09 mW/cm^2^), 488 nm 100% (3151.87 mW/cm^2^), 561 nm 100% (1332.86 mW/cm^2^), 594 nm 100% (801.48 mW/cm^2^) and 639 nm 100% (283.39 mW/cm^2^).

The stabilized step-function opsin (SSFO) channel variant, ChR2 C128S/D156A (Berndt et al., 2009, Bamann et al. 2010, Yizhar et al., 2011), has properties that make it attractive for a putative nanoscale focal activation approach; it is opened by blue-light and closed by green-light. This channel has been well-characterized by spectral analysis of photocurrent responses to different wavelengths, and has an optimal opening wavelength of 470, and optimal closing wavelength of 590 (Yizhar et al., 2011). We therefore used similar wavelengths to test coillumination. In single and co-illumination tests, however, we found maximal photocurrents of only 40 - 60 pA with full field activation of transfected neurons, as expected (**Fig. 4A, top**). In addition, 488 nm light activated channels were only partially closed with co-illumination of 561 nm (**Fig. 4A, top**) or 594 nm (**Fig. 4A, bottom**) light.

**Figure 4:**
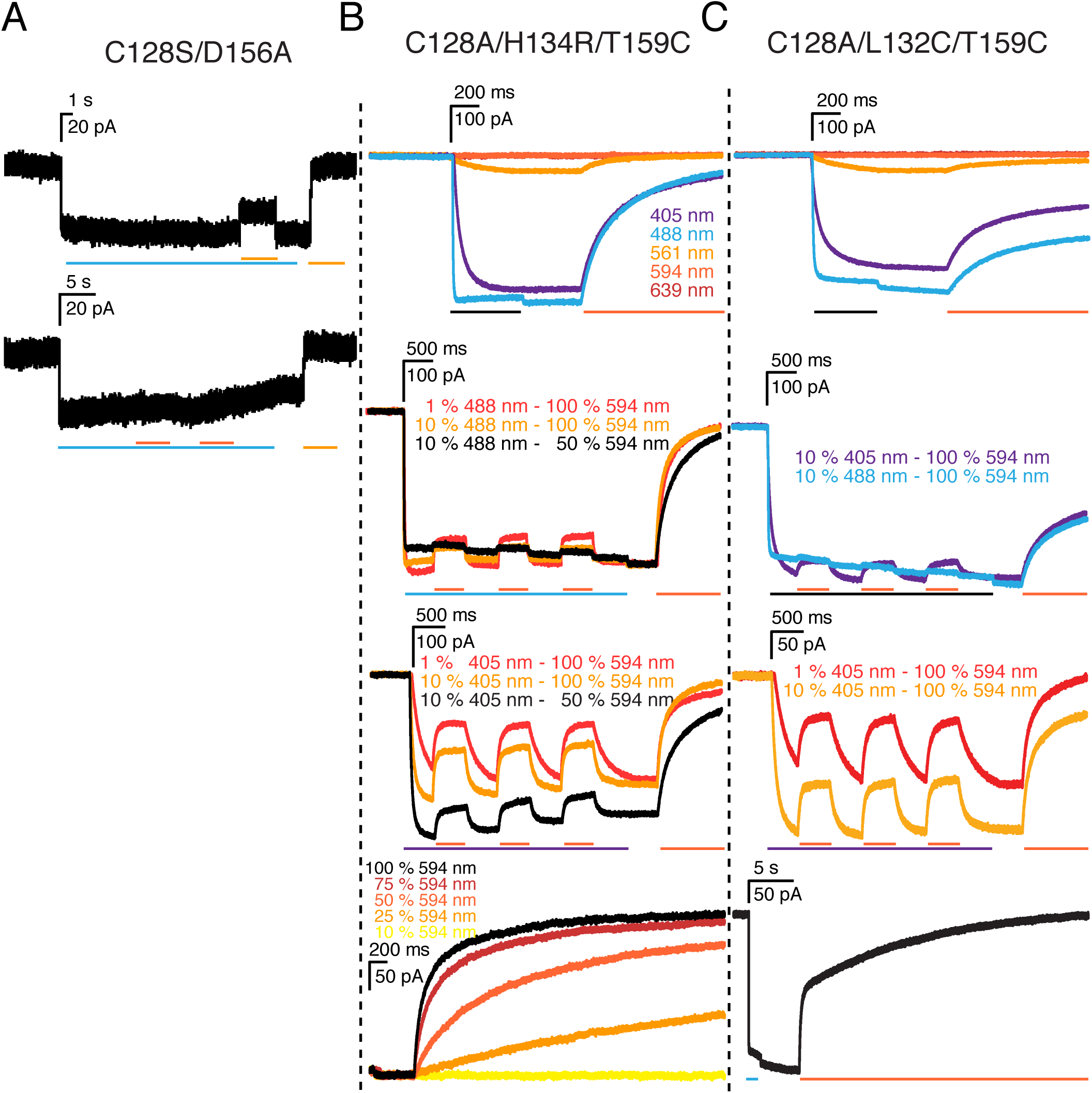
Illumination evoked responses of light-gated ion-channels with slow photocycles (step-function opsins). **A**) Responses to 405, 488, 561, 594 and 639 nm single wavelength laser illumination and coillumination of C128S/D156A (step-function opsin), C128/H134R/T159C (**B**), and C128S/L132C/T159C (**C**) switchable channels. Photocurrent recordings at specific wavelength illumination are indicated by the respective color of the recording. Scale bars above individual experiments indicate current (vertical) and time (horizontal). For coillumination experiments, bars below recordings indicate time of illumination for the indicated color-coded wavelength. Black bars refer to coillumination with the color of the depicted traces in cases where more than one coillumination wavelength was tested. Activating laser intensities were 405 nm 1% (22.5 mW/cm^2^) and 10% (86.83 mW/cm^2^), 488 nm 1% (79.66 mW/cm^2^) and 10% (295.6 mW/cm^2^), 561 nm 10% (146.3 mW/cm^2^), 594 nm 10% (80.1 mW/cm^2^) and 639 nm 10% (22.35 mW/cm^2^). For inactivation, coillumination intensites of 10%, 25%, 50%, 75% or 100% (801.48 mW/cm^2^) 594 nm light were used.

In an attempt to increase the photocurrent but maintain the switching capabilities of channels we made a ChR2 variant with C128A/H134R/T159C mutations. We reasoned that H134R would enhance stationary photocurrents (Nagel et al., 2005), T159C would increase photocurrents (Berndt et al., 2011), and C128A would allow the channel to be switched to a closed state with 561 nm illumination (and would inactivate more quickly than the C128S mutation (Berndt et al., 2009)). This channel exhibited photocurrents of 350 - 400 pA in response to 405 or 488 nm illumination and weak photocurrents when illuminated at 561 nm. The photocurrent was completely inactivated by single wavelength illumination with 594 nm light, and partially inactivated by coillumination. This channel was inactivated even more efficiently (with 594 nm light) when activated with 405 nm light (**Fig. 4B**). Thus, this channel appears more promising than the original SSFO channel for focal activation, in terms of larger photocurrent, and closing efficiency with inactivating light coillumination. We also generated a C128A/L132C/T159C channel, which should also have higher photocurrents and close with inactivating co-illumination, and in addition be permeable to calcium, thus allowing calcium imaging to detect activation. This channel did indeed have similar high photocurrents and inactivation properties compared to the C128A/H134R/T159C channel, although its closing kinetics were slower (**Fig. 4C**).

The C1V1 E122T/C167S channel is the corresponding step function version of the C1V1 channel. This channel maintained its red-shifted wavelength activation properties, and was activated by 488, 561, and 594 nm light, and weakly by 639 nm light. The C1V1 E122T/C167S channel showed efficient photocurrent reduction when stimulated with 561 nm light and coilluminated with 405 nm light, as indicated by earlier studies (Prigge et al., 2012); channels activated by 561 nm light could be efficiently closed by 405 nm light (but not by 488 nm light) (**Fig. 5A**). In contrast to the inactivating influence of 405 nm light on 561 nm activated currents, 405 nm coillumination following 488 nm light activation increased photocurrents further.

**Figure 5:**
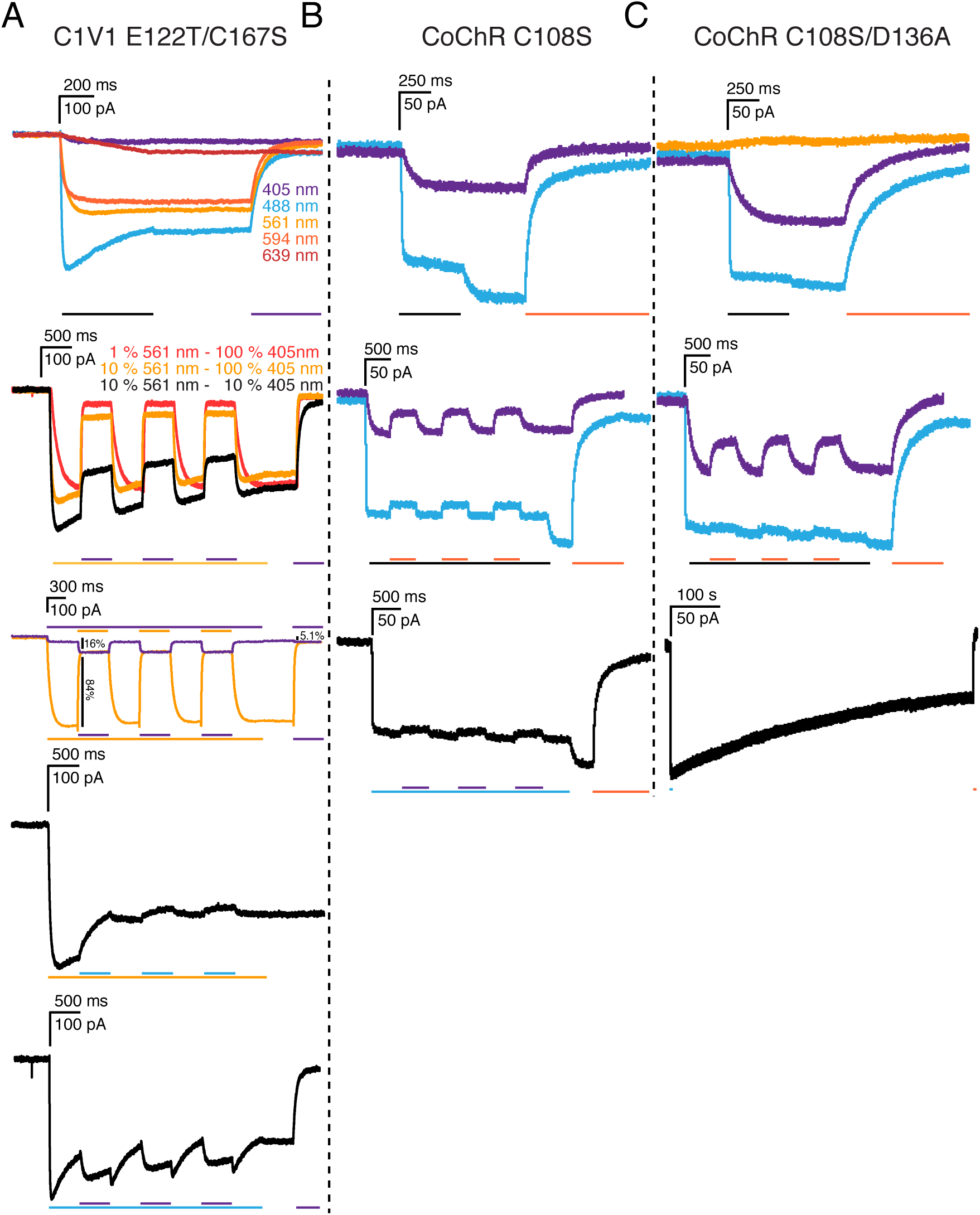
Illumination evoked responses of red-shifted and higher photocurrent light-gated ion-channels with step-function mutations. **A**) Responses to 405, 488, 561, 594 and 639 nm single wavelength laser illumination and coillumination of C1V1 E122T/C167S, CoChR C108S (B) and CoChR C108S/D136A (C) channels. Photocurrent recordings at specific wavelength illumination are indicated by the respective color of the recording. Scale bars above individual experiments indicate current (vertical) and time (horizontal). For coillumination experiments, bars below recordings indicate time of illumination for the indicated color-coded wavelength. Black bars refer to coillumination with the color of the depicted traces in cases where more than one coillumination wavelength was tested. Activating laser intensities were 405 nm 10% (86.83 mW/cm^2^), 488 nm 10% (295.6 mW/cm^2^), 561 nm 1% (42.05 mW/cm^2^ or 10% (146.3 mW/cm^2^), 594 nm 10% (80.1 mW/cm^2^) and 639 nm 10% (22.35 SmW/cm^2^). Inactivating laser intensities were 405 nm 10% (86.83 mW/cm^2^) and 100% (662.09 mW/cm^2^), 488 nm 100% (3151.87 mW/cm^2^), 561 nm 100% (1332.86mW/cm^2^), 594 nm 100% (801.48 mW/cm^2^) and 639 nm 100% (283.39 mW/cm^2^).

We further tested addition of the switchable mutations (C108S and D136A, corresponding to C128S and D156A in ChR2) to the CoChR channel variants, which showed the largest photocurrents of all channels tested. (Chronos variants showed intracellular accumulation in neurons resulting in small photocurrents and were therefore not tested further). The C108S mutation alone reduced the photocurrent of CoChR from ∼4 nA to 150 pA for 488 nm activation and to 50 pA for 405 nm activation, but did indeed cause the channel to be partially inactivated by 594 nm light, following activation with 405 or 488 nm light (**Fig. 5B**). In addition, the CoChR C108S mutant exhibited 488 nm mediated channel closure, as seen from the photocurrent increase after stopping illumination. Addition of the D136A mutation, to create CoChR C108S/D136A virtually eliminated this effect, and increased 405 nm evoked photocurrents, which were inactivated to a greater extent by 594 nm light (**Fig. 5C**).

As opposed to other approaches, the eNPAC construct - which expresses ChR2 H134R and NpHR3.0, a hyper-polarizing light-gated chloride pump, at equimolar amounts via connection by a P2A site that is cleaved intracellularly - does not depend on a specific illumination to close channels, but rather on a balance between blue light activation of ChR2 H134R to depolarize cells and red light activation of the NpHR3.0 proton pump to hyperpolarize cells (Gradinaru et al., 2010). Photocurrent responses from eNPAC-expressing neurons were detected for all tested wavelengths (**Fig. 6**). 405 and 488 nm evoked responses were almost identical to standard ChR2 H134R responses. However, 488 nm photocurrents displayed a small and brief transient after illumination ceased, most likely because NpHR3.0 is weakly activated by 488 nm light, and when this light is terminated the pumping of chloride-ions by NpHR3.0 stops more quickly than ChR2 H134R channels close, leading to a transient depolarization. This effect was not present for 405 nm excitation, implying that NpHR3.0 is not significantly activated by 405 nm light. When illuminated with 561 and 594 nm light, hyperpolarizing currents of ∼400 pA were recorded from eNPAC-expressing cells. Because ChR2 H134R is not activated by red wavelengths, these currents are predominantly from NpHR3.0 activation.

**Figure 6:**
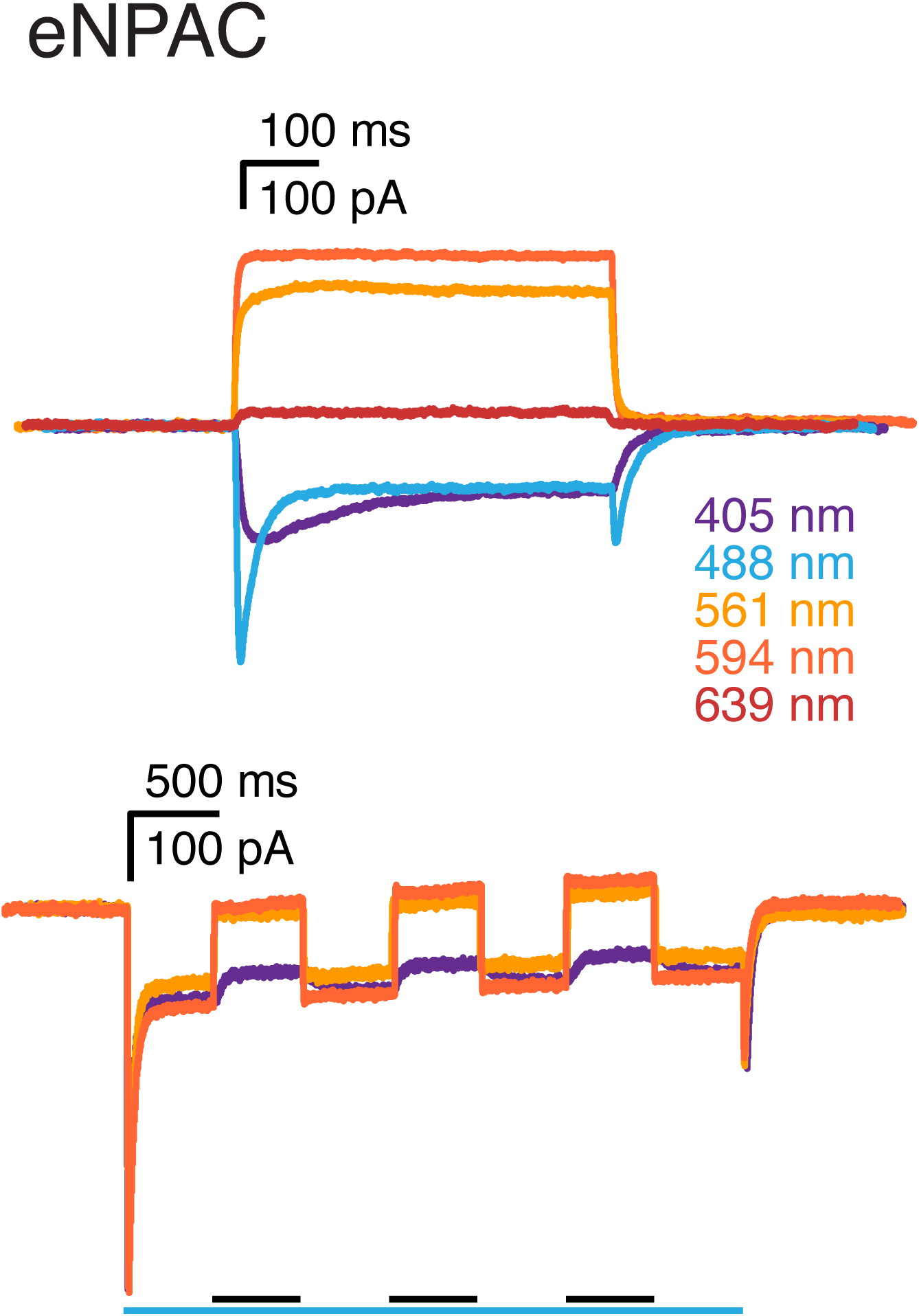
Illumination evoked responses of equimolar ChR2 H134R and NpHR3.0 (eNPAC). Evoked responses of eNPAC with 405, 488, 561, 594 and 639 nm single wavelength laser illumination (top) and coillumination (bottom). Scale bars above individual experiments indicate current (vertical) and time (horizontal). For coillumination experiments, bars below recordings indicate time of illumination for the indicated color-coded wavelength. Black bars refer to coillumination with the color of the depicted traces. Activating laser intensities were 10% (405 nm 86.83 mW/cm^2^; 488 nm 295.6 mW/cm^2^; 561 nm 146.3 mW/cm^2^) and 594 nm coillumination intensity was 100% (801.48 mW/cm^2^).

During initial channel tests we noted that even when 100% 594 nm laser power was used (801.48 mW/cm^2^), maximal channel inactivation was not reached. Therefore, we increased the 594 nm laser intensity by modifying the light path and exchanging the light guide from the DL594 laser module to the microscope with the beam splitter in the DirectFRAP unit (see Materials and Methods) and tested the effect of this modification on the most promising candidates activated by 594 nm light: ChR2 C128A/H134R/T159C and the CoChR C108S and C108S/D136 mutants (**Fig. 7**). ChR2 C128A/H134R/T159C channels were activated with 10% 488 nm laser power (41.92 W/cm^2^) or 20% 405 nm laser power (9.92 W/cm^2^ through the DirectFRAP laser path and inactivated by simultaneous illumination three consecutive times for 500 ms each, with 594 nm light of different intensities (**Fig. 7A**). The photocurrent values from these three exposures were pooled and compared to the average maximum photocurrent elicited by 500 ms single 488 or 405 nm illumination (between coillumination and a final 594 nm single wavelength exposure) to calculate % photocurrent reduction by coillumination with 594 nm light (**Fig. 7B**). Additionally, activation intensities from 1 to 20% were tested against 100% DL594 power (129.75 W/cm^2^) (**Fig. 7C**). Using the modified setup, we were able to reduce photocurrents of ChR2 C128A/H134R/T159C elicited with 488 nm light by 54%, and similar magnitude currents elicited with 405 nm light by 90%. Increased photocurrents were elicited by higher 488 nm activation intensities, which caused less efficient photocurrent reduction by coillumination. However, increased photocurrents elicited by higher (up to 20%) 405 nm activation were still efficiently reduced by coillumination.

**Figure 7:**
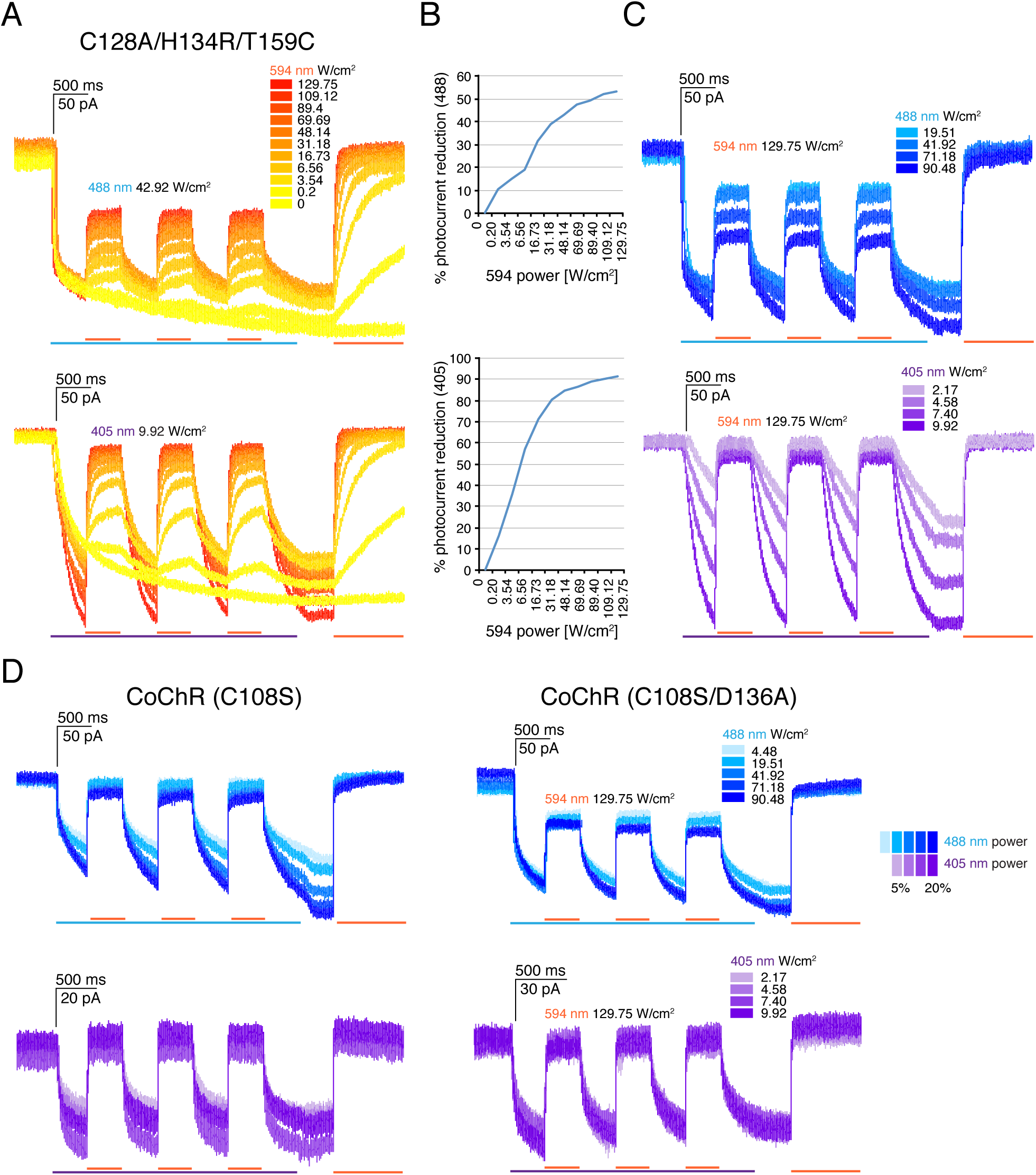
Reduction of 488 and 405 nm evoked photocurrents from ChR2 C128A/H134R/T159C, CoChR C108S and CoChR C108S/D136A channels. **A**) Effect of coillumination of different 594 nm laser powers on 10% 488 nm (top) or 20% 405 nm (bottom) evoked currents; 594 nm laser powers are indicated in the color-coded inset, and were tested in 10% steps from 0% to 100%. **B**) Relationship between 594 nm laser intensity and photocurrent reduction. **C**) Influence of activating 488 nm (top) or 405 nm (bottom) laser power (1%, 5%, 10% and 20%) on 594 nm light mediated photocurrent reduction. Scale bars indicate recorded currents (vertical) and time (horizontal). **D**) Photocurrent reduction of 488 nm light (top) and 405 nm light (bottom) evoked currents in CoChR C108S (left) and C108S/D136A (right) mutants. Trace and stimulation bar colors represent laser wavelengths and intensities (5%, 10%, 15%, 20%).

We then tested 488 and 405 nm activation of CoChR C108S and CoChR C108S/D136A channels at varying intensities with maximal 594 nm inactivation (**Fig. 7D**). Photocurrent reduction was most efficient for 405 nm activation and 594 nm coillumination. For CoChR C108S, photocurrents were higher for 488 nm than for 405 nm light activation, and 488 nm light evoked photocurrents inactivated only slightly less efficiently than those elicited by 405 nm light. CoChR C108S/D136A, on the other hand, had slightly higher photocurrents elicited by 405 nm light than 488 nm light, which were eliminated with 594 nm coillumination, while 488 nm elicited photocurrents were only inactivated by ∼50%. A photocurrent increase, when 488 nm illumination is discontinued was visible for the CoChR channel, and especially for the C108S mutant. In general, currents of the CoChR step function variants were weaker than the ChR2 step function variants and did not increase when higher 405 nm light intensities were used. However, 405 nm evoked currents could be efficiently reduced by exposure to 594 nm light. These experiments suggest that 405 nm light is the ideal activation wavelength for experiments where strong photocurrents and efficient photo-current reduction, using 594 nm coillumination, is needed.

To estimate the size of illuminated regions for which photocurrents can be detected, we tested focal single wavelength 488 nm light activation of proximal neuronal processes of cells expressing the ChR2 T159C channel. Photocurrents elicited by illumination of membrane areas with diameters down to 1.2 *μ*m could reliably be detected using electrophysiological patch-clamp recordings in the whole-cell configuration (**Fig. 8A**). However, detectable photocurrents were already weak, suggesting that photocurrents from smaller areas will be beyond detection limits, especially for switchable channel variants, which have reduced photocurrents. We therefore first investigated the feasibility of nanoscale optogenetic stimulation using a microscale approach and an illumination strategy borrowed from super resolution microscopy, in which a central area is illuminated with a wavelength of light that activates channelrhodopsins, while an overlapping donut region is simultaneously illuminated with a wavelength of light that inactivates these channels (**Fig. 8B**), thereby confining activation to a small central region. To accomplish this, we used photoactivation masks for 405 or 488 nm illumination of 16.7 *μ*m diameter areas, in combination with a donut-shaped 594 nm inactivation mask, resulting in a central activated region of 6.2 *μ*m. We first verified illumination regions using a fluorescently labelled slide (**Fig. 8C**). We then tested the approach on neurons transfected with eNPAC, a construct that co-expresses equimolar amounts of ChR2 H134R and NpHR3.0 (Gradinaru et al., 2010). In neurons transfected with eNPAC, coillumination with 594 nm light also effectively inactivated photocurrents evoked by 488 nm light (**Fig. 8D**). Stimulation of eNPAC expressing cells with 488 nm light in 6.2 or 16.7 *μ*m regions, resulted in ∼12 pA (**Fig. 8D, blue trace**) or ∼45 pA (**Fig. 8D, black trace**) steady-state photocurrents,respectively. When a 16.7 *μ*m spot of 488 nm light stimulation of ChR2 was combined with 594 nm costimulation of NpHR3.0 in an overlapping donut (resulting in a central activated region of approximately 6.2 *μ*m), net photocurrents could be tuned to be lower (**Fig. 8D**, **orange trace**), higher (**Fig. 8D**, **red trace**), or equivalent to (**Fig. 8D, black trace**) depolarization achieved with the 6.2 *μ*m stimulation mask alone. This demonstrates that NpHR3.0 currents are tunable in their strength by adjusting 594 nm laser power to counterbalance ChR2 mediated depolarization in the coilluminated area.

**Figure 8:**
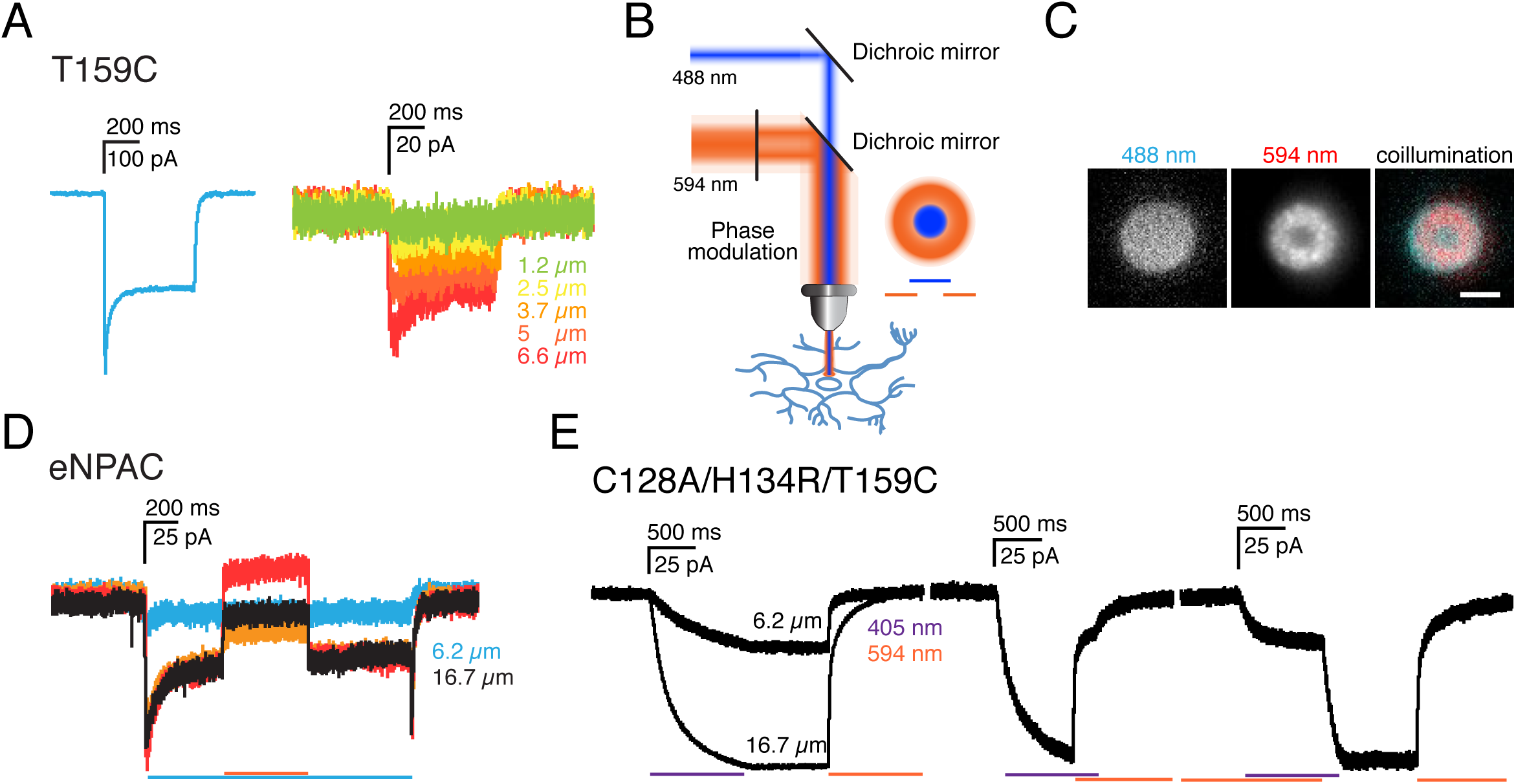
Electrophysiological recordings of photocurrents in focally stimulated areas. **A)** T159C channelrhodopsin photocurrents elicited by 20% 488 nm full field (left) or focal (right) activation of the cell body. **B**) Illumination scheme for focal activation based on STED/RESOLFT microscopy: 488 nm activation light is brought together with a 594 nm donut-shaped inactivation beam. The two wavelengths of illumination overlap and inactivate channels within the donut region, resulting in a potentially sub-diffraction limited central activated region. **C**) Images of yellow text highlighter, used as a dye for a uniform fluorescent field on a coverslip illuminated with 488 nm light in a 16.7 *μ*m diameter central region, and 594 nm light in an overlapping donut region, resulting in a central activated region of 6.2 *μ*m. Scale bar = 10 *μ*m. **D**) eNPAC photocurrents elicited by 488 nm light in a focal 6.2 *μ*m (blue) or 16.7 *μ*m (black) diameter region, or in a 16.7 *μ*m region combined with a 594 nm laser overlapping donut to inactivate the overlapping region resulting in a central activated region of 6.2 *μ*m (black trace). 594 nm laser intensity can be tuned to drive NpHR3.0-mediated hyperpolarizing currents to be weaker (orange trace), stronger (red trace) or equivalent to (black trace) the photocurrent elicited by 6.2 *μ*m stimulation of ChR2 (blue trace). **E**) Photocurrents of the ChR2 C128S/H134R/T159C variant evoked by 6.2 or 16.7 *μ*m regions of 20% 405 nm laser illumination and subsequent 594 nm illumination mediated closure (left), and with coillumination of 16.7 *μ*m 488 nm illuminated region and an overlapping donut of 594 nm illumination, resulting in a 6.2 *μ*m central activated region (right). When a 16.7 *μ*m 488 nm illuminated region was combined with a 594 nm overlapping donut region (leaving a central 6.2 *μ*m region of activation) the photocurrent is reduced to a level similar to that evoked by a 6.2 *μ*m region alone.

In a second approach the same illumination principle was applied to the ChR2 slow-photocyle variant C128A/H134R/T159C (**Fig. 8E**). The respective evoked photocurrents for 6.2 and 16.7 *μ*m wide areas, illuminated with 405 nm light, were 50 and 150 pA. When 16.7 *μ*m 405 nm photostimulation was combined with an overlapping donut of 594 nm illumination, photocurrents could be reduced to values close to those evoked by stimulation of a 6.2 *μ*m area alone, independent of the order in which the different wavelengths were applied (**Fig. 8E**).

## Discussion

We investigated a novel approach to control synaptic to sub-synaptic neuronal activity with optogenetics using switchable light-gated channels and an illumination approach inspired by STED/RESOLFT microscopy. Of all tested candidates, the ChR2 C128A/H134R/T159C and the slow CoChR variants C108S and C108S/D136A - activated with 405 nm light and inactivated by coillumination with 594 nm light - and the C1V1 E122T/C167S channel - activated by 561 nm light and inactivated by 405 nm light - were most promising in terms of highest photocurrents and efficient inactivation with coillumination. The combination of co-expression of a depolarizing cation-channel and hyperpolarizing light-gated chloride-pump (eNPAC) also represents an interesting alternative, in which counterbalancing currents can be used to cause a net voltage change of 0 mV in the area of 405 or 488, and 594 nm coillumination.

Based on our experiments, three approaches using light-gated ion-channels/pumps, could be used to achieve highly focal photostimulation. The first is simultaneous activating wavelength illumination of a central region, and inactivating wavelength coillumination in a donut overlapping this central activated region. The second (which could be used with step-function opsin channels that do not show sufficient inactivation during coillumination), is fast sequential illumination, in which channels are opened with one wavelength and subsequently closed in an outer donut region. For this approach, the time between initial channel opening and closing of the outer channels should be as short as possible. The third option makes use of the eNPAC construct to control depolarizing and hyperpolarizing currents with different wavelengths of light in central and surrounding regions. Theoretically the wavelengths can also be inverted, producing highly focal areas of hyperpolarization, mimicking inhibitory synapses.

Efficient inactivation of channels with coillumination usually required reducing activation laser intensities. It was reported that 594 nm inactivating light should be roughly 1000-fold more intense than 488 nm activating light to achieve photocurrent-inactivation of ∼95% for ChR2 C128S (Venkatachalam and Cohen, 2014). In our experiments, we tested the lowest available laser power for activation (1%; 4.48 W/cm^2^). The power density could be further reduced using filters or beam splitters, to achieve 1000-fold higher 594 nm power densities, however this would mainly reduce photocurrents from activated channels and not lead to a more efficient channel gating. Interestingly, channels were more efficiently inactivated by 594 nm light following 405 nm light activation, than following 488 nm light activation, even at only ∼100-fold higher 594 nm light power densities. We assume that 405 nm light illumination activates the P470 open state, but does so less efficiently. However, it is also possible that these channels have a separate conducting state for stimulation in the UV spectrum, which is more easily reduced by 594 nm illumination. In any case, 405 nm activation was superior to 488 nm activation, for subsequent inactivation by coilluminating 594 nm light to close channels. For eNPAC, 405 nm light evoked ChR2-mediated depolarization was completely counterbalanced by 80.15 - 160.3 mW/cm^2^ 594 nm illumination to activate hyperpolarizing NpHR3.0. Although these values fluctuated between experiments, they were minimal given approximate equimolar expression of ChR2 and NpHR3.0 in eNPAC. In this case, 405 nm activation light (rather than 488 nm) also avoids currents that arise from NpHR3.0 blue-light activation.

An important question is whether current optogenetic tools are powerful enough to allow detection at the nanoscale. Photocurrent recordings from *Xenopus* oocytes have yielded estimated single channel conductances of ∼50 fS for wild-type ChR2, with single channel currents of ∼5 fA (Nagel et al., 2003). Using non-stationary noise analysis, single channel Na^+^ conductances of wild-type ChR2 and ChR2 H134R in HEK cells were determined to be 41.5 fS at 200 mM extracellular Na^+^, with single channel currents of 3.5 fA at −60 mV (Feldbauer et al., 2009). Other studies using non-stationary noise analysis estimate ChR2 single channel conductance at around 1 pS, which principally would allow single channel recordings (Lin et al., 2009). However, the absence of such published recordings to date suggest this value may be an overestimation. During the stationary current photocycle phase, ∼60% of ChR2 H134R channels are expected to be in the open channel state (Feldbauer et al., 2009; Nagel et al., 2005). Thus, approximately 226 channels are active per *μ*m^2^ cell membrane in HEK cells (Feldbauer et al., 2009). If the proposed channel density in HEK cells translates to a similar presence in neuronal membranes, stimulation of a synapse size area (0.04 *μ*m^2^) would lead to the opening of ∼9 channels and currents of 31.5 - 45 fA at −60 mV. This is too low to be detected using standard electrophysiological methods. An increase in channel abundance in the membrane, e.g. by higher expression levels or targeting motifs to synapses, or increase in single channel conductance could enable detection of activation at the nanoscale. To mimic synaptic activation, ideally channels would possess properties that match currents elicited by AMPA receptors at excitatory synapses, where 1 - 2 AMPA receptors open during the peak of the postsynaptic current (Nimchinsky et al., 2004; Spruston et al., 1995). Single channel AMPA conductance is ∼8-10 pS (Jonas and Sakmann, 1992; Spruston et al., 1995). To match currents of a single AMPA receptor, the conductance of a wild-type ChR2 would need to be enhanced roughly 200-fold. Light-gated genetically-modified ionotropic glutamate channels (LiGluRs), which are based on AMPA receptors, and can be activated and inactivated by two different wavelengths of light, are another alternative (Berlin et al., 2016; Volgraf et al., 2006). LiGluR steady-state currents were reported to be ∼5-fold larger than those of wild-type ChR2 (Szobota et al., 2007), i.e. on the order of Chronos currents, but LiGluR requires the addition of an exogenous chemical photoswitch and may also alter the native firing properties of cells in which it is expressed, since it is based on the glutamate receptor that is natively present in the brain.

Electrophysiological readouts are currently the most sensitive method of detection of light-activated channels. But detection of focal activation by calcium or voltage sensors would verify focal activation. This would require that calcium or voltage sensors 1) can be imaged at wavelengths that do not influence light-gated channels, and 2) have a high enough signal-to-noise ratio to detect. GCaMP and other green calcium indicators could in principle be used in combination with red-shifted channelrhodopsins, but at present, there are no red-shifted channels that are not activated in the blue spectrum at wavelengths necessary to excite GCaMP and other green calcium dyes. Another option is to use a red calcium indicator, like RCaMP (Akerboom et al., 2013), with ChR2-based channels, which are only activated by blue light. Although RCaMP-based calcium indicators have lagged behind the superior signal-to-noise ratio of GCaMP-based calcium indicators, which would be required for detection of focal activation at the nanoscale, newly generated red calcium sensors appear more promising (Dana et al., 2016). Although the signal-to-noise ratio of voltage-sensing dyes is often not as high as for calcium indicators, and they possess a substantial spectral overlap with ChR2, recent Archaerhodopsin-based voltage indicators reportedly allow detection of miniature excitatory postsynaptic depolarizations using near infrared light excitation (Hochbaum et al., 2014).

Although we demonstrate the feasibility of a focal optogenetics approach at the microscale, we also show that at present channel conductance is too low to allow nanoscale activation. Nevertheless, this work may be useful for future development of nanoscale optogenetics. It is likely that stronger conducting channel variants will be found. New single channel optogenetic tools with faster switching kinetics that allow depolarization and hyperpolarization would allow nanoscale activation, however. For instance, the ion selectivity of ChR2 has recently been selectively modified to create a chloride-conduction channel (Berndt et al., 2014; Wietek et al., 2014). It would also be interesting to test the sdChR C138S/E154A variant, because it has been reported to possess good photocurrent inhibition (Venkatachalam and Cohen, 2014). In addition, optogenetics now extends beyond light-gated channels. Inducing protein conformational changes or interactions in nanodomains by light, for example, may be feasible using this technique.

## Materials&Methods

### Animals

Use of animals for experimentation was approved and performed according to the specifications of the Institutional Animal Care and Ethics Committees of Goettingen University (T10.31) and of the German animal welfare laws.

### Mammalian expression constructs

Vectors for CaMKII α-controlled expression of channelrhodopsins (pAAV-CaMKII α-hChR2-mCherry/EYFP-WPRE) including the H134R, T159C, E123T/T159C and C128S/D156A ChR2 variants (Berndt et al., 2011; Yizhar et al., 2011), ChR1 VChR1 chimera C1V1 E122T/E162T (Erbguth et al., 2012), and the eNpHR3.0-EYFP (halorhodopsin) and hChR2(H134R)-mCherry fusion construct eNPAC (pAAV-hsyn-eNPAC) (Gradinaru et al., 2010), are available from Karl Deisseroth (Stanford University/Howard Hughes Medical Institute). The L132C channelrhodopsin (CatCh) variant (Kleinlogel et al., 2011), was provided by Ernst Bamberg (Max-Planck-Institute for Biophysics, Germany). With the assistance of GenScript, pAAV-CaMKII α-hChR2(H134R)-mCherry-WPRE, pAAV-CaMKII α-hChR2(T159C)-mCherry-WPREand pAAV-CaMKII α-C1V1(E122T/E162T)-TS-mCherry were mutated to combine common ChR characteristics and yield the slow-photocycle switchable high-conducting variants hChR2(C128A/H134R/T159C), hChR2(C128A/L132C/T159C) and C1V1(E122T/C167S) in the same backbone. Additional ChRs selected for their strong photocurrents, Chronos and CoChR, together with their mutated slow-photocycle switchable variants, Chronos C145S and C145S/E162T and CoChR C108S and C108S/D136A, under CMV promotor control, are available from Edward Boyden (Massachusetts Institute of Technology).

### Dissociated primary hippocampal cultures

Dissociated hippocampal cultures were cultured on 25 mm coverslips (Paul Marienfeld GmbH; cat. no. 0117650) coated with 0.04% polyethylenimine (PEI) (Sigma; cat. no. P3143) in 6-well plates (CytoOne; cat. no. CC7672-7506); 2 ml of 0.04% PEI in dH_2_O, stored at −20 °C, was added per well at room temperature for approximately 14 hours, followed by 3 washes with dH_2_O after which plate were directly used for cultures or stored in dH_2_O at 4 °C until needed. The hippocampal culture method was adapted from Gary Banker (Goslin and Banker, 1991; Kaech and Banker, 2006). Pregnant Wistar rats were sacrificed between E18 and E19 by CO_2_. Embryos were extracted and brains transferred to 10 cm dishes containing dissection medium (Hank’s Balanced Salt Solution + 10 mM HEPES). Hemispheres were separated, meninges removed, and hippocampi extracted and collected in a 15 ml falcon-tube containing dissection medium, on ice. Tissue was digested in 2 ml 0.25% Trypsin-EDTA (Gibco; cat. no. 25200-056) for 20 min at 37 °C, washed 3 times with 5 ml dissection medium, triturated in 1 ml plating medium (Dulbecco’s Modified Eagle Medium with 10% FBS; Gibco), and filtered through a 100 *μ*m cell strainer (Corning, Inc.; cat. no. 352360). Cell yield was determined using trypan blue staining (Sigma; cat. no. T8154), and cells plated at densities of 30,500 to 37,000 cells/cm^2^ per coverslip and cultured at 37 °C and 5% CO_2_. The following day plating medium was exchanged for 2 ml culture medium (Neurobasal with 1X B-27 supplement, 1X Glutamax and Penicillin (5000 U/ml)/Streptomycin (5000 *μ*g/ml); all from Gibco). 5 *μ*M AraC was added once on DIV7.

### Calcium Phosphate Transfection

On DIV3, conditioned culture medium was collected and exchanged for pre-equilibrated (37 °C and 5% CO_2_) Opti-MEM (Gibco; cat. no. 11058-021). 1/3^rd^ fresh culture medium was added to the collected conditioned medium which was sterile filtered through a 0.22 *μ*m syringe-driven filter (Millipore; cat. no. SLGV033RS) and stored in the incubator. For each well of a 6-well plate, 7.5 *μ*l 2 M CaCl_2_ and 3.5 - 4 *μ*g plasmid DNA in 60 *μ*l dH_2_O were mixed and 60 *μ*l transfection buffer (274 mM NaCl, 10 mM KCl, 1.4 mM Na_2_HPO_4_, 15 mM glucose, 42 mM Hepes, pH 7.05-7.12) was then added drop-wise, under gentle agitation. This solution was incubated for 20 min at room temperature and then gently added to cells which were returned to the incubator for 90 min. The Opti-MEM plus transfection mix is then exchanged for 37 °C and 10% CO_2_ pre-equilibrated Neurobasal medium, returned to the incubator for 10 min, and subsequently exchanged for the stored conditioned medium.

### STED imaging

Hippocampal neurons transfected with ChR2-EYFP were imaged using a custom-built STED microscope. Excitation pulses of 490 nm wavelength were delivered using a pulsed-laser diode (Toptica Photonics, Graefelfing, Germany), followed by pulses at 595 nm delivered by a Ti:Sapphire laser (MaiTai; Spectra-Physics, Darmstadt, Germany) for stimulated depletion with a donut shaped focal profile created by introducing a polymeric phase plate (RPC Photonics, Rochester, NY) into the path of the STED beam. The STED and excitation pulse beams were synchronized by external triggering, overlapped with a custom-made dichroic mirror, and focused into a 1.3 NA objective lens (PL APO, CORR CS, 63×, Leica, Wetzlar, Germany). Fluorescence was passed through a 535/50 band-pass filter, and collected with an optical fiber connected to an avalanche photodiode (PerkinElmer, Waltham, MA). Images were recorded with resonant mirror scanning along the x-axis and stage scanning along the y-axis at 10 *μ*s pixel dwell times.

### FRAP experiments

Transfected hippocampal neurons at DIV12 growing on coverslips were transferred into a low-profile RC-41LP imaging chamber (Warner Instruments; cat. no. 64-0368) filled with 1 ml extracellular solution (in mM: 145.4 NaCl, 5 KCl, 2 CaCl_2_, 1 MgCl_2_, 10 HEPES, 7 glucose in dH_2_O; 300 mOsm, pH 7.4). Live imaging was conducted on a Zeiss Axio Observer.Z1 microscope equipped with a 405 nm 50 mW diode laser, 488 nm 100 mW OPSL, 561 nm 40 mW diode laser and 639 nm 30 mW diode laser, a TIRF slider module, a laser manipulation DirectFRAP module and an Evolve 512 EMCCD camera (Photometrics). Image acquisition was controlled with AxioVision 4.8.2 software (Zeiss). Images were collected using a 100x objective (alpha Plan-Apochromat 100x/1.46 oil DIC (UV) Vis-IR) and laser intensities of 10% and 1 s intervals with a gain of 800 and exposure times below 1 s. Following stable baseline acquisition for at least 10 s, defined regions of cell bodies and processes were bleached for 1 s with 100% 488 nm laser intensity using the Laser Manipulation with DirectFRAP tool and photoactivation mask with 2.5 *μ*m diameter. Average intensities were recorded from the FRAP region and 3 additional regions of interest: two different control regions on the cell and one off the cell to detect background noise. Background noise was subtracted, signal from control areas were used to compensate for bleaching and data was normalized by setting the average of fluorescence values from 1 to 10 s before bleaching at 100%. Tau (τ), was computed for bleach recovery curves fit with a single exponential curve using Prism v5.0d software (GraphPad).

### Electrophysiology and Optogenetics

Electrophysiological recordings were made from DIV8-DIV12 hippocampal neurons transfected with light-gated channels/pumps on DIV3. Cultures were transferred to a custom-made imaging chamber, adapted to fit with the motorized stage, in extracellular solution (in mM: 148 NaCl, 2.4 KCl, 2 CaCl_2_, 1 MgCl_2_, 10 HEPES, 7 glucose in dH_2_O; 300 mOsm, pH 7.4). To isolate ChR2 photocurrents and block spontaneous network activity, 1 *μ*M TTX (Alomone Labs; cat. no. T-550), 50 ΩM APV (Abcam; cat. no. ab120003), 10 *μ*M CNQX (Sigma; cat. no. C239) and 10 ΩM gabazine (Abcam; cat. no. ab120042) was added to the extracellular solution. Patch pipettes with 3 - 5 MΩ resistance were made from fire polished borosilicate capillaries (Harvard Apparatus; cat. no. 300060, OD 1.5 mm x ID 0.86 mm) with a P-97 Micropipette Puller (Sutter Instruments). Silver wire electrodes were chlorinated with a 2 M KCl solution using an ACl-01 apparatus (npi electronic). Patch pipettes were backfilled with 7 *μ*l internal solution (in mM: 130 potassium gluconate, 8 KCl, 2 CaCl_2_, 1 MgCl_2_, 10 EGTA, 10 HEPES, 2 Mg-ATP, 0.3 GTP-Na; 290 - 295 mOsm, pH 7.3), using 20 *μ*l Microloader pipette tips (Eppendorf; cat. no. 5242 956.003). A pipette holder was controlled using a MPC-385-2 micromanipulator system (Sutter Instruments). Whole-cell patch-clamp recordings were obtained with an EPC10 USB double patch-clamp amplifier and Patchmaster software (HEKA) at 25 kHz sampling intervals.

All optogenetic experiments were conducted on an inverted Zeiss Axio Observer.Z1 microscope equipped with a 405 nm 50 mW diode laser, 488 nm 100 mW OPSL, 561 nm 40 mW diode laser and 639 nm 30 mW diode laser, 20x LD A-Plan, 40x and 63x EC Plan-NEOFLUAR and 100x α Plan-APOCHROMAT objectives, a TIRF slider module, a laser manipulation DirectFRAP module and an Evolve 512 EMCCD camera (Photometrics). An additional 594 nm 100 mW DPSS laser (Rapp OptoElectronic GmbH; Hamburg, Germany) was added to the epi-fluoresence light path with a 940 *μ*m light guide, together with custom photomasks, complementing the existing DirectFRAP photomasks by adding a 594 nm border (2 *μ*m for 100x objective) around the DirectFRAP photomasks. A photomask of a 150 *μ*m wide area or no photomask was used for initial single wavelength full-field illumination tests. All lasers were operated by TTL pulses delivered by the HEKA amplifier.

Laser powers were determined with a laser power meter (Fieldmate) and a silicon OP-2 VIS optical sensor (Coherent GmbH; Dieburg, Germany) at the back aperture of the 40x and 100x objectives, using two aluminum masks to represent the respective exit pupil diameter, 10.7 and 4.8 mm. There was no difference in the measured power between those two masks. Laser intensities were measured after 4 - 6 hours of use and with the TIRF/FRAP beamsplitter set at 50% TIRF/50% FRAP. The measured laser powers were subsequently corrected for the respective transmission of the objective. Laser output generally fluctuated ±30 *μ*W and the measurement error was previously determined to be ∼10%. 594 nm laser intensity was measured using the 200 *μ*m light guide and respective powers for the 100 *μ*m light guide were extrapolated. For initial tests of light-gated channel candidates, we used wide-field illumination with the 40x objective at power densities of 405 nm 86.83 mW/cm^2^, 488 nm 295.6 mW/cm^2^, 561 nm 146.3 mW/cm^2^ and 594 nm 801.48 mW/cm^2^ or 1.08 W/cm^2^. In experiments in which the light path was adjusted to increase 594 nm laser power (by exchanging the 940 *μ*m light guide for a 100 *μ*m light guide, and the DirectFRAP beamsplitter for an AHF beam splittter), power densities using the 40x objective were 405 nm 27.10 - 64.56 mW/cm^2^ 488 nm 228.1 mW/cm^2^, 561 nm 133.4 mW/cm^2^, 594 nm 129.75 W/cm^2^ or 267.25 W/cm^2^. Because the 594 nm photomasks did not cover the whole wide-field illuminated area, FRAP photomasks of similar area were used for stimulation of 405 nm light at 9.92 W/cm^2^ and 488 nm light at 41.92 W/cm^2^.

Because channels with a stable open state are sensitive to broad-spectrum daylight conditions, channels were closed before and after each experiment by a brief illumination with the respective closing wavelength. Channelrhodopsins were characterized by their responses to 500 ms light pulses of 405, 488, 561, 594 and 639 nm, using the DL594 laser and no photomask for the 594 nm and the TIRF laser for the other wavelengths. To test focal optogenetic stimulation and record responses, 488 nm light pulses were delivered using DirectFRAP photomasks and the 100x objective; 594 nm Rapp Opto custom-designed photomasks were overlaid to produce a co-illumination donut surrounding a central region.

Microscope and components including lasers were controlled with Zeiss AxioVision and Zen Blue software. Transfected cells were identified by fluorescence and selected based on health and membrane integrity. Recordings were acquired from cells voltage-clamped at −70 mV. Liquid junction potential, pipette and cell capacitance influences were compensated. Recordings were managed with IGOR Pro (Wavemetrics; version 6.22A) and the Patcher’s Power Tools extension for HEKA files, provided by the Department of Membrane Biophysics at the Max Planck Institute for Biophysical Chemistry in Goettingen. In cases where small shifts in timing of laser pulses occurred, traces were manually aligned to illumination onset for comparison.

## Acknowledgements

We thank Stefan Hell for insightful discussions and the use of a STED setup for imaging channelrhodopsins, and a custom-built RESOLFT setup to test nanoscale stimulation of neurons expressing channelrhodopsins. This work was supported by a Sofja Kovalevskaja grant from the Alexander von Humboldt Foundation, and European Research Council starting grant SytActivity FP7 260916 to C.D., and the Deutsche Forschungsgemeinschaft funded Center for Nanoscale Microscopy and Molecular Physiology of the Brain (CNMPB).

## References

Akerboom, J., Carreras Calderon, N., Tian, L., Wabnig, S., Prigge, M., Tolo, J., et al. (2013). Genetically encoded calcium indicators for multi-color neural activity imaging and combination with optogenetics. Front Mol Neurosci 6, 2. doi:10.3389/fnmol.2013.00002.

Bamann, C., Gueta, R., Kleinlogel, S., Nagel, G., and Bamberg, E. (2010). Structural guidance of the photocycle of channelrhodopsin-2 by an interhelical hydrogen bond. Biochemistry 49, 267–278. doi:10.1021/bi901634p.

Bamann, C., Kirsch, T., Nagel, G., and Bamberg, E. (2008). Spectral characteristics of the photocycle of channelrhodopsin-2 and its implication for channel function. J Mol Biol 375, 686–694. doi:10.1016/j.jmb.2007.10.072.

Berlin, S., Szobota, S., Reiner, A., Carroll, E. C., Kienzler, M. A., Guyon, A., et al. (2016). A family of photoswitchable NMDA receptors. Elife 5. doi:10.7554/eLife.12040.

Berndt, A., Lee, S. Y., Ramakrishnan, C., and Deisseroth, K. (2014). Structure-guided transformation of channelrhodopsin into a light-activated chloride channel. Science (80-.). 344, 420–424. doi:10.1126/science.1252367.

Berndt, A., Yizhar, O., Gunaydin, L. A., Hegemann, P., and Deisseroth, K. (2009). Bi-stable neural state switches. Nat Neurosci 12, 229–234. doi:10.1038/nn.2247.

Dana, H., Mohar, B., Sun, Y., Narayan, S., Gordus, A., Hasseman, J. P., et al. (2016). Sensitive red protein calcium indicators for imaging neural activity. Elife 5. doi:10.7554/eLife.12727.

Deisseroth, K. (2011). Optogenetics. Nat Methods 8, 26–29. doi:10.1038/nmeth.f.324.

Feldbauer, K., Zimmermann, D., Pintschovius, V., Spitz, J., Bamann, C., and Bamberg, E. (2009). Channelrhodopsin-2 is a leaky proton pump. Proc Natl Acad Sci U S A 106, 12317–12322. doi:10.1073/pnas.0905852106.

Gradinaru, V., Zhang, F., Ramakrishnan, C., Mattis, J., Prakash, R., Diester, I., et al. (2010). Molecular and cellular approaches for diversifying and extending optogenetics. Cell 141, 154–165. doi:10.1016/j.cell.2010.02.037.

Gunaydin, L. A., Yizhar, O., Berndt, A., Sohal, V. S., Deisseroth, K., and Hegemann, P. (2010). Ultrafast optogenetic control. Nat Neurosci 13, 387–392. doi:10.1038/nn.2495.

Hochbaum, D. R., Zhao, Y., Farhi, S. L., Klapoetke, N., Werley, C. A., Kapoor, V., et al. (2014). All-optical electrophysiology in mammalian neurons using engineered microbial rhodopsins. Nat Methods 11, 825–833. doi:10.1038/nmeth.3000.

Jonas, P., and Sakmann, B. (1992). Glutamate receptor channels in isolated patches from CA1 and CA3 pyramidal cells of rat hippocampal slices. J Physiol 455, 143–171. Available at: http://www.ncbi.nlm.nih.gov/pubmed/1282929.

Klapoetke, N. C., Murata, Y., Kim, S. S., Pulver, S. R., Birdsey-Benson, A., Cho, Y. K., et al. (2014). Independent optical excitation of distinct neural populations. Nat. Methods 11, 338–346. doi:10.1038/nmeth.2836.

Kleinlogel, S., Feldbauer, K., Dempski, R. E., Fotis, H., Wood, P. G., Bamann, C., et al. (2011). Ultra light-sensitive and fast neuronal activation with the Ca(2)+-permeable channelrhodopsin CatCh. Nat Neurosci 14, 513–518. doi:10.1038/nn.2776.

Lin, J. Y., Knutsen, P. M., Muller, A., Kleinfeld, D., and Tsien, R. Y. (2013). ReaChR: a red-shifted variant of channelrhodopsin enables deep transcranial optogenetic excitation. Nat Neurosci. doi:10.1038/nn.3502.

Lin, J. Y., Lin, M. Z., Steinbach, P., and Tsien, R. Y. (2009). Characterization of engineered channelrhodopsin variants with improved properties and kinetics. Biophys J 96, 1803–1814. doi:10.1016/j.bpj.2008.11.034.

Mohanty, S. K., Reinscheid, R. K., Liu, X., Okamura, N., Krasieva, T. B., and Berns, M. W. (2008). In-depth activation of channelrhodopsin 2-sensitized excitable cells with high spatial resolution using two-photon excitation with a near-infrared laser microbeam. Biophys J 95, 3916–3926. doi:10.1529/biophysj.108.130187.

Nagel, G., Brauner, M., Liewald, J. F., Adeishvili, N., Bamberg, E., and Gottschalk, A. (2005). Light activation of channelrhodopsin-2 in excitable cells of Caenorhabditis elegans triggers rapid behavioral responses. Curr Biol 15, 2279–2284. doi:10.1016/j.cub.2005.11.032.

Nagel, G., Szellas, T., Huhn, W., Kateriya, S., Adeishvili, N., Berthold, P., et al. (2003). Channelrhodopsin-2, a directly light-gated cation-selective membrane channel. Proc Natl Acad Sci U S A 100, 13940–13945. doi:10.1073/pnas.1936192100.

Nikolic, K., Grossman, N., Grubb, M. S., Burrone, J., Toumazou, C., and Degenaar, P. (2009). Photocycles of channelrhodopsin-2. Photochem Photobiol 85, 400–411. doi:10.1111/j.1751-1097.2008.00460.x.

Nimchinsky, E. A., Yasuda, R., Oertner, T. G., and Svoboda, K. (2004). The number of glutamate receptors opened by synaptic stimulation in single hippocampal spines. J Neurosci 24, 2054–2064. doi:10.1523/JNEUROSCI.5066-03.2004.

Packer, A. M., Peterka, D. S., Hirtz, J. J., Prakash, R., Deisseroth, K., and Yuste, R. (2012). Two-photon optogenetics of dendritic spines and neural circuits. Nat Methods 9, 1202–1205. doi:10.1038/nmeth.2249.

Papagiakoumou, E., Anselmi, F., Begue, A., de Sars, V., Gluckstad, J., Isacoff, E. Y., et al. (2010). Scanless two-photon excitation of channelrhodopsin-2. Nat Methods 7, 848–854. doi:10.1038/nmeth.1505.

Prigge, M., Schneider, F., Tsunoda, S. P., Shilyansky, C., Wietek, J., Deisseroth, K., et al. (2012). Color-tuned Channelrhodopsins for Multiwavelength Optogenetics. J Biol Chem. doi:10.1074/jbc.M112.391185.

Schikorski, T., and Stevens, C. F. (1997). Quantitative ultrastructural analysis of hippocampal excitatory synapses. J Neurosci 17, 5858–5867. Available at: http://www.ncbi.nlm.nih.gov/pubmed/9221783.

Schoenenberger, P., Grunditz, A., Rose, T., and Oertner, T. G. (2008). Optimizing the spatial resolution of Channelrhodopsin-2 activation. Brain Cell Biol 36, 119–127. doi:10.1007/s11068-008-9025-8.

Spruston, N., Jonas, P., and Sakmann, B. (1995). Dendritic glutamate receptor channels in rat hippocampal CA3 and CA1 pyramidal neurons. J. Physiol. 482 (Pt 2), 325–52. Available at: http://www.ncbi.nlm.nih.gov/pubmed/7536248 [Accessed June 4, 2018].

Stehfest, K., Ritter, E., Berndt, A., Bartl, F., and Hegemann, P. (2010). The branched photocycle of the slow-cycling channelrhodopsin-2 mutant C128T. J Mol Biol 398, 690–702. doi:10.1016/j.jmb.2010.03.031.

Szobota, S., Gorostiza, P., Del Bene, F., Wyart, C., Fortin, D. L., Kolstad, K. D., et al. (2007). Remote Control of Neuronal Activity with a Light-Gated Glutamate Receptor. Neuron 54, 535–545. doi:10.1016/J.NEURON.2007.05.010.

Venkatachalam, V., and Cohen, A. E. (2014). Imaging GFP-based reporters in neurons with multiwavelength optogenetic control. Biophys J 107, 1554–1563. doi:10.1016/j.bpj.2014.08.020.

Volgraf, M., Gorostiza, P., Numano, R., Kramer, R. H., Isacoff, E. Y., and Trauner, D. (2006). Allosteric control of an ionotropic glutamate receptor with an optical switch. Nat Chem Biol 2, 47–52. doi:10.1038/nchembio756.

Wietek, J., Wiegert, J. S., Adeishvili, N., Schneider, F., Watanabe, H., Tsunoda, S. P., et al. (2014). Conversion of channelrhodopsin into a light-gated chloride channel. Science (80-.). 344, 409–412. doi:10.1126/science.1249375.

Yizhar, O., Fenno, L. E., Prigge, M., Schneider, F., Davidson, T. J., O’Shea, D. J., et al. (2011). Neocortical excitation/inhibition balance in information processing and social dysfunction. Nature 477, 171–178. doi:10.1038/nature10360.

Zhang, Y. P., and Oertner, T. G. (2007). Optical induction of synaptic plasticity using a light-sensitive channel. Nat Methods 4, 139–141. doi:10.1038/nmeth988.

